# Molecular and phytochemical investigation of medicinal *Rhododendron* spp. in the Qinghai–Xizang Plateau: the botanical sources of the traditional Tibetan medicine “Tale”

**DOI:** 10.64898/2026.11.05.764495

**Authors:** Qundong Liu, Siyuan Xie, Xiaoxue Huang, Xue Liu, Yongsheng Wei, Zhiwei Zhang, Xiaomei Zhang, Weihan Qin, Xiang Liu

**Affiliations:** Chongqing Key Laboratory for the Utilization and Evaluation of Special Chinese Materia Medica Resources, Chongqing Academy of Chinese Materia Medica, Chongqing, China; Daba Mountain Medicinal Animals and Plants Chongqing Observation and Research Station, Chengkou, China; Key Lab. of Tibetan Medicine Resources Conservation and Utilization of Tibet Autonomous Region, Tibet Agriculture & Animal Husbandry University, Nyingchi, China

**Keywords:** *Rhododendron*, *psb*A–*trn*H, molecular identification, flavonoid, active compounds, traditional Tibetan medicine

## Abstract

The Qinghai–Xizang Plateau in China is a hotspot for *Rhododendron* species, with approximately 400 species distributed across the region. Various *Rhododendron* species in this area have been used medicinally in traditional Tibetan medicine for a long time and are collectively referred to as “Tale.” Furthermore, “Tale” is classified as “Talegabao” or “Talenabao,” depending on the species. These two types of medicine have similar but distinct properties, and “Talenabao” is usually used as an alternative to “Talegabao.” However, differences in their phytochemical compositions and methods for discriminating between them have not been examined. To solve these problems limited the use of “Tale” medicines, 46 plant specimens belonging to nine *Rhododendron* species were collected from the Qinghai–Xizang Plateau. Molecular methods based on the internal transcribed spacer 2 (ITS2) region of nuclear DNA and *psb*A–*trn*H regions of chloroplast DNA were applied to these samples. In addition, 10 flavonoid compounds were quantitatively determined by HPLC. The results indicated *psb*A–*trn*H region successfully distinguished the species used as “Talegabao” or “Talenabao,” whereas ITS2 failed to distinguish them because of its high degree of conservation. All 10 compounds were detected in all samples. However, the species used as “Talegabao” had more than twice the total content of the 10 flavonoid compounds observed in the species used as “Talenabao.” Moreover, differences in the contents of certain compounds among species were detected, indicating potential chemical markers for the quality evaluation of “Tale” medicines. “Talegabao” is the first choice among “Tale” medicines. “Talenabao” can be used as an alternative to “Talegabao”; however, the appropriate dose requires further discussion in practical use. The *psb*A–*trn*H region is recommended as a DNA barcode for distinguishing between these two types of medicines.

## Introduction

The genus *Rhododendron* (Ericaceae) comprises approximately 1,000 species distributed widely throughout the Northern Hemisphere, particularly in Asia [1]. *Rhododendron* species are known not only for their ornamental flowers but also for their traditional medicinal uses in many countries [2]. The Qinghai–Xizang Plateau in China is notable for its rich and diverse flora, with approximately 400 *Rhododendron* species distributed throughout the region, making it a major center of *Rhododendron* diversity [3]. Owing to their wide distribution and species richness in this area, various *Rhododendron* species have a long history of medicinal use in traditional Chinese, Tibetan, and Mongolian medicine, among others [4,5].

In traditional Tibetan medicine, *Rhododendron* species play an important role, with more than 80 species used medicinally. These medicines are called “Dama” (སྟག་མ) or “Tale” (ད་ལིས།), also known as “Dali,” “Tali,” or “Balu” [4,6]. “Dama” consists of the dried flowers, leaves, and seeds of certain evergreen species with large leaves, including *R. przewalskii* Maxim., *R. arboreum* var. *roseum* Lindl., *R. pingianum* Fang, *R. coryanum* Tagg & Forrest, *R. aganniphum* Balf. f. & Kingdon-Ward, *R. phaeochrysum* var. *agglutinatum* (Balf. f. & Forrest) D. F. Chamb., and *R. principis* var. *vellereum* Hutch. ex Tagg, which mostly belong to Section *Ponticum*. “Tale” consists of the dried flowers, leaves, and twigs of certain evergreen species with small leaves and scales on the surface of leaves and twigs. In practice, “Tale” is classified as “White Tale” and “Black Tale.” “White Tale,” locally called “Talegabao” (ད་ལིས་དཀར་པོ།), is characterized by plants with greenish-yellow upper leaf surfaces and yellowish or light-brown lower surfaces, grayish-white branches and twigs, and white, yellowish, or light-pink corollas. More than 17 species are used as “Talegabao,” including *R. anthopogonoides* Maxim., *R. primuliflorum* Bur. & Franch., *R. cephalanthum* Franch., *R. hypenanthum* Balf.f., *R. anthopogon* D.Don, *R. mainlingense* S.H.Huang & R.C.Fang, and *R. micranthum* Turcz., which mostly belong to Section *Pogonanthum*. “Black Tale,” locally called “Talenabao” (ད་ལིས་ནག་པོ།), is characterized by plants with dark-green or gray-green leaves, gray to black branches and twigs, and red, purple, or bluish-violet corollas. More than 20 species are used as “Talenabao,” including *R. capitatum* Maxim., *R. nivale* Hook.f., *R. thymifolium* Maxim., *R. intricatum* Franch., and *R. nitidulum* Rehder & E.H. Wilson, which mostly belong to the *Rhododendron* subsection *Lapponica* [4].

In clinical practice, the two types of “Tale” medicines have similar but somewhat distinct effects. “Talegabao” is described as having effects that clear heat and reduce inflammation; relieve cough and asthma; strengthen the stomach and body, combat aging, and treat disorders caused by rLung (wind), chibpa (bile), and pechen (phlegm), as well as lung diseases, laryngitis with hoarseness, asthma attributed to environmental adaptation issues, bronchitis, emphysema, spleen and stomach deficiency with coldness, indigestion, gastric ptosis, gastric dilation, gastric cancer, hepatosplenomegaly, and edema. It is also applied externally to reduce inflammation and swelling. “Talenabao” is described as having effects that clear heat and reduce inflammation, relieve cough and asthma, strengthen the stomach, disperse swelling, nourish the kidneys and enhance vitality, and combat aging; it is also used to treat lung pain and laryngitis [4]. In practice, local medical workers consider “Talegabao” to have a better effect and prefer to collect it, particularly *R. anthopogonoides* and *R. primuliflorum*, which are usually the first choice. “Talenabao” is usually used as an alternative.

Multiple flavonoid compounds have been isolated from *R. anthopogonoides*, including quercetin, isoquercetin, avicularin, and dihydroquercetin [7]. In addition, owing to the strong scent of these plants, compounds in their essential oils have attracted attention [8]. Pharmacological studies have shown that *R. anthopogonoides* extracts have expectorant, antitussive, antiasthmatic [4], bactericidal [9], anti-inflammatory [10], smooth muscle–relaxing, and sedative effects. In addition, the essential oil has shown insecticidal activity, suggesting its potential as a biological insecticide [11,12]. However, few studies have reported the chemical composition of other species used as “Tale.” The chemical basis of “Tale” medicines and the differences between “Talegabao” and “Talenabao” remain unclear. Owing to the broad geographic distribution of traditional Tibetan medicine and the use of multiple *Rhododendron* species, different regions may prefer different species. For example, both *R. anthopogonoides* and *R. primuliflorum* are used as “Talegabao”, however, *R. anthopogonoides* mostly distributed in Qinghai, Gansu provinces and *R. primuliflorum* distributed in Xizang, Yunnan and Sichuan provinces, which leading to confusion regarding the botanical sources of the drugs (Fig 1). Moreover, standards for “Tale” medicines are lacking. Accurate identification and quality assessment have not been adequately established. The wide distribution and morphological similarities of *Rhododendron* species also complicate their classification, with the taxonomic status of some species remaining debated and nomenclature varying among regions. This situation can lead to confusion regarding the use of “Tale” crude drugs.

**Fig 1.**
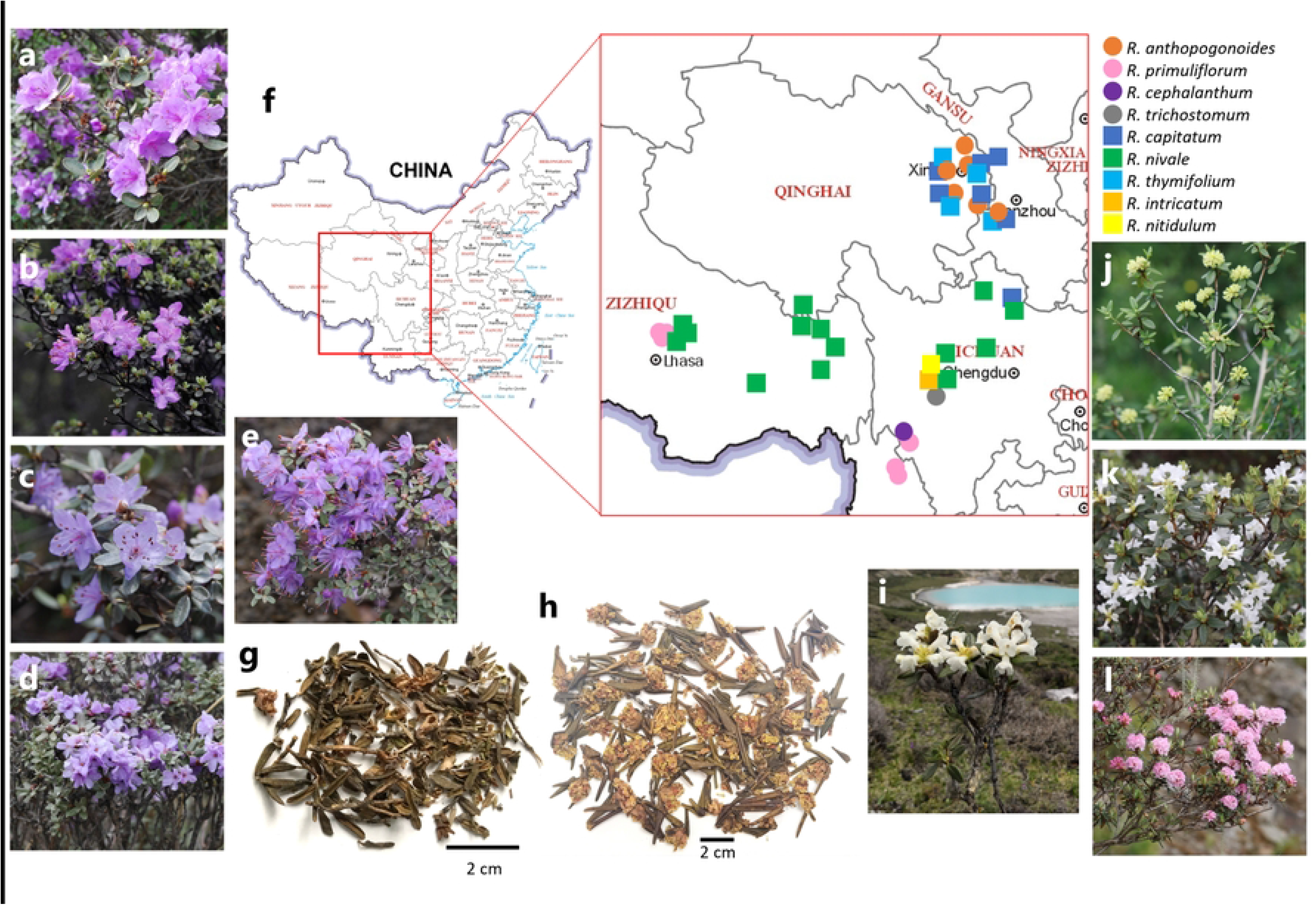
*Rhododendron* plant specimens and “Tale” medicines. a, *R. capitatum*; b, *R. nivale*; c, *R. thymifolium*; d, *R. intricatum*; e, *R. nitidulum*; f, locations of *Rhododendron* plant specimens; g, medicines of “Talenabao”; h, medicines of “Talegabao”; i, *R. trichostomum*; j, *R. anthopogonoides*; k, *R. primuliflorum*; l, *R. cephalanthum*.

*Rhododendron* is one of the most taxonomically challenging and species-rich genera for species identification and phylogenetic inference. Molecular methods based on nuclear and chloroplast DNA have been evaluated for discriminating among *Rhododendron* species. Liu et al. used the internal transcribed spacer 2 (ITS2) region to classify 10 *Rhododendron* species and found that ITS2 sequences successfully discriminated among them [13]. Yan et al. evaluated four DNA barcodes using 531 samples representing 173 *Rhododendron* species. Among the four barcodes, the *psb*A–*trn*H region provided the highest species resolution [14]. These findings provided a basis for developing molecular methods to identify *Rhododendron* species used as “Tale” medicines.

In this study, we aimed to discriminate among different “Tale” species and clarify the phytochemical differences between “Tale” medicines. Forty-six plant specimens and nine crude drug samples belonging to nine medicinally used *Rhododendron* species were collected from the Qinghai–Xizang Plateau for investigation. Molecular methods based on the ITS2 region of nuclear DNA and the *psb*A–*trn*H region of chloroplast DNA were applied to the plant specimens. In addition, 10 flavonoid compounds, including dihydromyricetin, hyperoside, isoquercitrin, taxifolin, guaijaverin, avicularin, astragalin, quercitrin, quercetin, and naringenin, were quantified using high-performance liquid chromatography (HPLC). Variations in compound composition were characterized using multivariate statistical analysis, specifically orthogonal partial least squares discriminant analysis (OPLS-DA).

## Materials and Methods

### Plant specimens and drug samples

Corolla, leaves, and twigs of 46 plant specimens representing nine *Rhododendron* species, including *R. anthopogonoides*, *R. primuliflorum*, *R. nivale*, *R. thymifolium*, *R. capitatum*, *R. trichostomum*, *R. cephalanthum*, *R. intricatum*, and *R. nitidulum*, were collected from the Qinghai–Xizang Plateau, China (Table 1; Fig 1). In addition, nine crude drug samples were purchased from drug markets in Lhasa, Chengdu, and Xining, China (Table 2). The plant specimens and crude drug samples were morphologically authenticated by the corresponding author. All samples were sealed in plastic bags and stored at room temperature (15°C–25°C) away from sunlight. Voucher specimens were deposited at the Chongqing Academy of Chinese Materia Medica in Chongqing, China.

**Table 1.** Plant specimens of *Rhododendron*.

| Section | Scientific name | Voucher No. | Collected from |
|---|---|---|---|
| *Rhododendron* Sect. *Pogonanthum* (“Talegabao”) | *R. anthopogonoides* | RP1 | Beishan, Huzhu, Qinghai |
|  |  | RP2 | Beishan, Huzhu, Qinghai |
|  |  | RP3 | Shierpan, Yuanfu, Bazha, Huzhu, Qinghai |
|  |  | RP4 | Wanglvdaban, Xianmi, Menyuan, Qinghai |
|  |  | RP5 | Qunjialinchang, Huangzhong, Qinghai |
|  |  | RP6 | Jianglalinchang, Guide, Qinghai |
|  |  | RP7 | Dalijiashan, Xunhua, Qinghai |
|  |  | RP8 | Linxia, Gansu |
|  | *R. primuliflorum* | RP9 | Gaba, Tanggu, Linzhou, Xizang |
|  |  | RP10 | Pangduo, Linzhou, Xizang |
|  |  | RP11 | Songpan, Linzhou, Xizang |
|  |  | RP12 | Linzhou, Xizang |
|  |  | RP13 | Habaxueshan, Sanba, Xianggelila, Yunnan |
|  |  | RP14 | Pujinnima, Deqin, Yunnan |
|  |  | RP15 | Chitu, Daocheng, Sichuan |
|  |  | RP16 | Yading airport, Daocheng, Sichuan |
|  | *R. cephalanthum* | RP17 | Echushan, Geka, Daocheng, Sichuan |
|  | *R. trichostomum* | RP18 | Zheduoshan, Kangding, Sichuan |
| *Rhododendron* Sect. *Rhododendron* Subsect. *Lapponica* (“Talenabao”) | *R. capitatum* | RL1 | Garang, Guide, Qinghai |
|  |  | RL2 | Jianglalinchang, Guide, Qinghai |
|  |  | RL3 | Beishan, Huzhu, Qinghai |
|  |  | RL4 | Qunjialinchang, Huangzhong, Qinghai |
|  |  | RL5 | Dalijiashan, Xunhua, Qinghai |
|  |  | RL6 | Linxia, Qinghai |
|  |  | RL7 | Wutailing, Tianzhu, Gansu |
|  |  | RL8 | Xueshanliang, Songpan, Sichuan |
|  | *R. nivale* | RL9 | Youxi, Jitang, chaya, Changdu, Xizang |
|  |  | RL10 | Basu, Changdu, Xizang |
|  |  | RL11 | Leiwuqi, Changdu, Xizang |
|  |  | RL12 | Shangka, Leiwuqi, Changdu, Xizang |
|  |  | RL13 | Qialashan, Songpan, Linzhou, Lhasa, Xizang |
|  |  | RL14 | Baga, Wumatang, Dangxiong, Lhasa, Xizang |
|  |  | RL15 | Lagen, Gongtang, Dangxiong, Lhasa, Xizang |
|  |  | RL16 | Sejilashan, Nyingchi, Xizang |
|  |  | RL17 | Huya, Pingwu, Sichuan |
|  |  | RL18 | Xinduqiao, Kangding, Sichuan |
|  |  | RL19 | Gedaliangzi, Danba, Sichuan |
|  |  | RL20 | Balangshan, Xiaojin, Sichuan |
|  |  | RL21 | Rangkou, Hongyuan, Sichuan |
|  |  | RL22 | Nangqian, Yushu, Qinghai |
|  | *R. thymifolium* | RL23 | Beishan, Huzhu, Qinghai |
|  |  | RL24 | Garang, Guide, Qinghai |
|  |  | RL25 | Qunjialinchang, Huangzhong, Qinghai |
|  |  | RL26 | Linxia, Gansu |
|  | *R. intricatum* | RL27 | Xinduqiao, Kangding, Sichuan |
|  | *R. nitidulum* | RL28 | Bamei, Daofu, Sichuan |

**Table 2.** Crude drug samples of “Tale”.

| Local name | Derived from | Voucher No. | Market |
|---|---|---|---|
| Talegabao | *R. anthopogonoides* | D1 | Chengdu, Sichuan |
|  |  | D2 | Xining, Qinghai |
|  |  | D3 | Xining, Qinghai |
|  |  | D4 | Xizang |
|  |  | D5 | Xizang |
|  | *R. primulaeflorum* | D6 | Xizang |
|  |  | D7 | Lhasa, Xizang |
|  |  | D8 | Xizang |
|  |  | D9 | Zhangshu Market, Jiangxi |

### Isolation of total DNA

Total DNA was extracted from 30 mg of dried leaf samples using a Plant DNA Isolation Kit (FOREGENE Co., Ltd., Chengdu, China) according to the manufacturer’s instructions, with some modifications. A 1-μL aliquot of the extraction solution from each sample was examined using 1.0% agarose gel electrophoresis.

### PCR amplification and sequence analysis of ITS2 and *psb*A-–*trn*H regions

Thirty-eight plant specimens representing nine species were selected for ITS2 sequence analysis. All 46 plant specimens representing the nine species were subjected to *psb*A–*trn*H sequence analysis. The ITS2 and *psb*A–*trn*H regions were amplified by PCR using two pairs of primers: ITS2_F (5′-ATG CGA TAC TTG GTG TGA AT-3′), ITS2_R (5′-GAC GCT TCT CCA GAC TAC AAT-3′), and *psb*A–*trn*H_F (5′-GTT ATG CAT GAA CGT AAT GCT C-3′), and *psb*A–*trn*H_R (5′-CGC GCG TGAT GAA TTC ACA ATC C-3′), respectively. A 25-μL PCR mixture contained 2 μL of total DNA as the template, 1 μL of 10 μM each primer, and 12.5 μL of 2× PCR HeroTM Mix (FOREGENE Co., Ltd., Chengdu, China), which contained Foregene Taq Hero, dNTPs, Tris-HCl (pH 8.8), KCl, MgCl2, and DNA loading buffer. PCR amplification was performed using an Arhat 96 PCR Thermal Cycler (Monad Biotech Co., Ltd., Suzhou, China). The cycling conditions included an initial hot start at 94°C for 3 min, followed by 35 cycles of denaturation at 94°C for 10 s, annealing at 55°C for 10 s, and extension at 72°C for 20 s, with a final extension at 72°C for 5 min. A 1-μL aliquot of each PCR product was examined using 1.0% agarose gel electrophoresis. The PCR products were then bidirectionally sequenced by Tsingke Biotech Co., Ltd. (Chongqing, China) using the primers employed for the PCR amplification. Sequencing data were assembled using BioEdit (version 7.0.0), and primer sequences were removed using MEGA (version 10.1.6). Consensus sequences were aligned and compared using Multalin software (http://multalin.toulouse.inra.fr/multalin/) or BioEdit (version 7.0.0).

### Chemicals and reagents

Chemical standards of quercitrin, isoquercitrin, hyperoside, quercetin, dihydromyricetin, guaijaverin, taxifolin, astragalin, avicularin, and naringenin were purchased from Beijing Bethealth People Biomedical Technology Co., Ltd. (Beijing, China), with purity ≥98%. HPLC-grade methanol and acetonitrile were purchased from TEDIA (Fairfield, OH, USA), whereas HPLC-grade phosphoric acid was procured from Aladdin (Shanghai, China). Analytical-grade methanol was sourced from Chongqing Chuandong Chemical (Group) Co., Ltd. (Chongqing, China). Ultrapure water (18.2 MΩ·cm) was obtained using a Milli-Q Integral 5 water purification system (Millipore, Billerica, MA, USA).

### Preparation of standard and sample solutions for HPLC analysis

Standard solutions were prepared using the corresponding reference compounds. Quercetin, isoquercetin, hyperoside, quercetin, dihydromyricetin, guaijaverin, taxifolin, astragalin, avicularin, and naringenin (2.79, 2.69, 2.59, 2.55, 2.57, 2.50, 2.69, 2.87, 2.78, and 2.67 mg, respectively) were accurately weighed and dissolved in 25 mL of methanol. Subsequently, 1 mL of each standard solution was mixed to obtain concentrations of 11.16, 10.76, 10.36, 10.20, 10.28, 10.00, 10.76, 11.48, 11.12, and 10.68 μg·mL^−1^ for each compound, respectively. The standard mixture solution was further diluted 2-, 4-, 5-, 8-, and 10-fold. Calibration curves were generated by plotting the peak area against the corresponding concentration of each analyte.

*Rhododendron* specimens and crude drug samples were heated at approximately 50°C until dry, ground into powder, and passed through a 65-mesh (250-μm) sieve. The resulting fine powder was stored at room temperature away from sunlight until analysis. Subsequently, 500 mg of fine powder from each sample and 50 mL of methanol were added to a covered conical flask. The flask was weighed before extraction and then extracted in a water bath at 70°C for 60 min. After the solution was cooled to room temperature, the flask was weighed again, and methanol was added to restore the original weight. The supernatant was collected after static settling and filtered through a 0.22-μm filter (Tianjin Jinteng Experiment Equipment Co., Ltd., Tianjin, China) for HPLC analysis. Each sample was prepared in triplicate.

### HPLC conditions

The HPLC system (Agilent, Santa Clara, CA, USA) was equipped with a G7111A quaternary pump, G1322A degasser, G7129A autosampler, G1316A column oven, and G7115A DAD detector. HPLC analysis was conducted using an Agilent 1260 system (Agilent, USA) with a Welch UltimateXB-C18 column (4.6 mm × 250 mm, 5 μm). The mobile phase consisted of solvent A (acetonitrile) and solvent B (0.1% phosphoric acid in water, v/v), with the following gradient elution program: 0–5 min, 15% A; 5–25 min, 30% A; 25–45 min, 45% A; 45–60 min, 60% A; 60–87 min, 80% A; 87–90 min, 10% A; and 90–100 min, 10% A. The flow rate was 1.0 mL/min, and the column temperature was maintained at 30°C. Detection was performed at 254 nm with a 10 μL injection volume.

### HPLC method validation

Precision was evaluated by performing six analyses of a standard solution containing 10 standard compounds, and the RSD of the peak area for each compound was calculated. Repeatability was assessed by analyzing six sample solutions, with variations expressed as RSD. The stability of one sample solution was evaluated by analyzing it at 0, 4, 8, 12, and 24 h. The accuracy of the method was assessed using a recovery test. The standard compounds were weighed to approximately 100% of their original amounts and added to the RL13 sample powder. The spike sample was then extracted and analyzed using the aforementioned method.

### Data processing

Independent two-sample t-tests or one-way analysis of variance were conducted using SPSS Statistics (version 20; IBM, USA). The data matrix used for OPLS-DA was generated from the HPLC peak areas of the 55 samples, and the data were normalized using the Z-score transformation method, as described in our previous study [15]. SIMCA-P (version 14.1; Umetrics, Umea, Sweden) was used to perform the OPLS-DA.

## Results

### Molecular identification of *Rhododendron* species

Thirty-eight ITS2 sequences and 46 *psb*A–*trn*H sequences from nine species were successfully amplified and determined. The 46 *psb*A–*trn*H sequences were all 379 bp in length and showed high similarity, with several single-nucleotide differences detected among them. All sequences were classified into six sequence types (Table 3). Among the species in Section *Pogonanthum* (“Talegabao”), eight *R. anthopogonoides* specimens showed identical sequences. *R. cephalanthum* and *R. trichostomum* showed the same sequence, with guanine at alignment position 242, whereas eight *R. primuliflorum* specimens showed identical sequences, with thymine at position 187 and guanine at position 242. However, sequences from the “Talenabao” species in Section *Rhododendron*, Subsection *Lapponica*, including seven *R. capitatum*, 12 *R. nivale*, four *R. thymifolium*, one *R. intricatum*, and one *R. nitidulum*, showed the same sequence, with five differences at alignment positions 229 (G), 235 (G), 242 (G), 255 (A), and 262 (G) compared to the sequence of *R. anthopogonoides*. In addition, two *R. nivale* sequences (RL13 and RL16) had cytosine at position 105, and one *R. capitatum* sequence (RL7) had thymine at positions 255 and 262. The sequence alignment results indicated that the *psb*A–*trn*H region successfully classified the species used as “Talegabao” and “Talenabao.” Furthermore, among the four species used as “Talegabao,” the *psb*A–*trn*H region clearly distinguished *R. anthopogonoides* and *R. primuliflorum* from *R. cephalanthum* and *R. trichostomum*.

**Table 3.** *psb*A–*trn*H and ITS2 sequences detected in *Rhododendron* species.

|  | Local name | Scientific name | Voucher No. | 105 | 187 | 229 | 235 | 242 | 255 | 262 | Acession Number |
|---|---|---|---|---|---|---|---|---|---|---|---|
| *psb*A-*trn*H region | Talegabao | *R. anthopogonoides* | RP1~8 | T | C | A | C | T | T | T | PZ896252 |
|  |  | *R. cephalanthum, R. trichostomum* | RP17, RP18 | * | * | * | * | G | * | * | PZ896253 |
|  |  | *R. primuliflorum* | RP9~16 | * | T | * | * | G | * | * | PZ896254 |
|  | Talenabao | *R. capitatum, R. nivale, R. thymifolium, R. intricatum, R. nitidulum* | RL1~6, RL8~12, RL14~15,RL17~28 | * | * | G | G | G | A | G | PZ896255 |
|  |  | *R. nivale* | RL13, RL16 | C | * | G | G | G | A | G | PZ896256 |
|  |  | *R. capitatum* | RL7 | * | * | G | G | G | * | * | PZ896257 |
| ITS2 region |  |  |  | 36 | 40 | 65 | 84 | 132 | 236 | 266 | Acession Number |
|  | Talegabao & Talenabao | *R. anthopogonoides, R. primuliflorum, R. cephalanthum, R. trichostomum, R. capitatum, R. nivale, R. intricatum, R. nitidulum* | RP4, RP10, RP13~18; RL1~7, RL9~10, RL12~19, RL27~28 | T | T | C | G | C | C | C | PZ888515 |
|  | Talegabao | *R. anthopogonoides* | RP1 | A | A | * | * | * | * | * | PZ888518 |
|  |  | *R. anthopogonoides* | RP7 | * | A | * | * | * | * | * | PZ888516 |
|  | Talenabao | *R. capitatum*, *R. nivale* | RL8, 11 | * | * | * | * | * | A | * | PZ888520 |
|  |  | *R. nivale* | RL20 | * | * | * | * | * | * | T | PZ888522 |
|  |  | *R. nivale* | RL21 | * | * | T | * | * | A | * | PZ888519 |
|  |  | *R. nivale* | RL22 | * | * | * | A | * | * | * | PZ888517 |
|  |  | *R. thymifolium* | RL23~26 | * | * | * | * | T | * | T | PZ888521 |

Numbers above sequence indicate the alignment positions. Asterisk (\*) indicates nucleotide is same with that in the first sequence.

Compared with *psb*A–*trn*H sequences, ITS2 sequences were more conserved. All 38 ITS2 sequences were 428 bp in length. Among them, 27 sequences from 8 species, excluding *R. thymifolium*, were identical (Table 3). Several nucleotide differences were detected among the other seven sequences from *R. anthopogonoides*, *R. capitatum*, and *R. nivale*, indicating intraspecific nucleotide variation in individual plants of these species. The four *R. thymifolium* specimens had identical sequences. Two positions, 132 and 262, contained thymine and distinguished this species from others. In particular, thymine at position 132 was detected only in *R. thymifolium*, indicating that this position may be species-specific.

### Quantitative analysis of the 10 compounds in “Tale”

The quantitative method was validated by evaluating linearity (R2), precision, stability, repeatability, and recovery (Table 4). High correlation coefficients were observed for all 10 analytes (R2 ≥ 0.9990), indicating strong correlation between concentration and peak area within the tested range (1.000–11.48 μg·mL^−1^). Precision, repeatability, and stability were also validated for each analyte, with relative standard deviations (RSDs) ranging from 0.96% to 2.57%, 1.48% to 2.97%, and 0.25% to 2.91%, respectively. Recoveries of the 10 analytes ranged from 94.43% to 106.08%, with RSDs ranging from 1.66% to 2.89%. These findings indicate that the established method is sufficiently precise and accurate for quantitative analysis of the 10 compounds in *Rhododendron*.

**Table 4.** Regression equations, correlation coefficients, linearity ranges, precisions, repeatability, stability, and recovery for the 10 analytes.

| Compound | Calibration Curve Equation | R<sup>2</sup> | Linear Range/µg·mL<sup>-1</sup> | Precision (RSD%) n=6 | Stability (RSD%) n=6 | Repeatability (RSD%) n=6 | Recovery (%) n=6 | RSD% |
|---|---|---|---|---|---|---|---|---|
| Dihydromyricetin | Y= 4.3625X-0.4832 | 0.9996 | 1.028 ~ 10.28 | 2.57 | 2.91 | 2.44 | 104.54 | 2.89 |
| Hyperoside | Y=26.7X-2.1122 | 0.9998 | 1.036 ~ 10.36 | 1.06 | 0.25 | 2.97 | 94.93 | 2.01 |
| lsoquercitrin | Y=26.562X+1.7001 | 0.9998 | 1.076 ~ 10.76 | 1.04 | 0.88 | 2.24 | 106.08 | 2.75 |
| Taxifolin | Y=3.1571X+0.5621 | 0.9990 | 1.076 ~ 10.76 | 0.81 | 2.21 | 2.48 | 104.66 | 1.69 |
| Guaijaverin | Y=23.52X+1.1895 | 0.9998 | 1.000 ~ 10.00 | 0.96 | 0.41 | 2.11 | 95.34 | 1.66 |
| Avicularin | Y=26.846X+1.1688 | 0.9998 | 1.112 ~ 11.12 | 0.98 | 0.18 | 1.48 | 100.47 | 2.19 |
| Astragalin | Y=17.884X+2.285 | 0.9997 | 1.148 ~ 11.48 | 0.98 | 0.92 | 2.01 | 100.86 | 2.53 |
| Quercitrin | Y=28.144X+1.9873 | 0.9998 | 1.116 ~ 11.16 | 0.97 | 1.3 | 2.35 | 95.50 | 2.74 |
| Quercetin | Y=40.472X-2.2232 | 0.9998 | 1.020 ~ 10.20 | 0.97 | 0.57 | 2.28 | 103.58 | 2.68 |
| Naringenin | Y=5.8574X+0.4178 | 0.9999 | 1.068 ~ 10.68 | 0.97 | 0.64 | 2.95 | 100.42 | 2.73 |

Ten compounds were simultaneously detected in the HPLC chromatograms of all plant specimens and crude drug samples based on their retention times. The relative contents of the 10 compounds were also determined. The contents of all 10 compounds varied among the plant specimens and crude drug samples (Table 5, Fig 2). Dihydromyricetin, hyperoside, isoquercitrin, taxifolin, guaijaverin, avicularin, astragalin, quercitrin, quercetin, and naringenin had contents ranging from 0.163 to 1.106, 0.052 to 4.523, 0.036 to 3.704, 0.273 to 3.214, 0.058 to 0.704, 0.023 to 1.232, 0.030 to 1.643, 0.030 to 2.697, 0.057 to 1.524, and 0.106 to 1.610 mg·g^−1^, respectively. The total content of the 10 compounds ranged from 1.220 to 3.836 mg·g^−1^, corresponding to 0.24%–2.24% of the samples. Clear differences were detected between “Talegabao” and “Talenabao.” In the “Talegabao” samples, the 10 compounds—dihydromyricetin, hyperoside, isoquercitrin, taxifolin, guaijaverin, avicularin, astragalin, quercitrin, quercetin, and naringenin—had average contents of 0.332, 2.016, 0.929, 1.665, 0.386, 0.326, 0.459, 1.029, 0.743, and 0.454 mg·g^−1^, respectively. The average total content was 8.340 mg·g^−1^, with an average percentage of 1.65%. In contrast, the compounds in “Talenabao” had relatively low contents. Dihydromyricetin, hyperoside, isoquercitrin, taxifolin, guaijaverin, avicularin, astragalin, quercitrin, quercetin, and naringenin had average contents of 0.409, 0.723, 0.266, 0.801, 0.196, 0.192, 0.277, 0.306, 0.519, and 0.323 mg·g^−1^, respectively. The average total content was 4.011 mg·g^−1^, with an average percentage of 0.80%, which was less than half that of “Talegabao.” The 10 detected compounds differed in content between the two types of medicines. Except for dihydromyricetin, which had a lower content in “Talegabao,” the other nine compounds had higher contents in “Talegabao” than in “Talenabao.” In particular, hyperoside, isoquercitrin, taxifolin, guaijaverin, quercitrin, and quercetin had significantly higher contents in “Talegabao,” with all p-values < 0.001, indicating that these six compounds contributed substantially to the difference in total content between the two types of medicine. Among the “Talegabao” samples, the four species *R. anthopogonoides*, *R. primuliflorum*, *R. cephalanthum*, and *R. trichostomum* showed similar total contents of the 10 compounds, with average values of 8.411, 8.227, 8.948, and 8.167 mg·g^−1^, respectively. Several compounds differed in content among species. *R. anthopogonoides* had significantly lower contents of dihydromyricetin and quercitrin than *R. primuliflorum*, with p-values of 0.0019 and 0.013, respectively. *R. anthopogonoides* had a significantly higher quercetin content than *R. primuliflorum*, with a p-value of 0.01. In addition, *R. cephalanthum* had high contents of taxifolin and naringenin and low levels of avicularin and astragalin. *R. trichostomum* had a high content of avicularin and low contents of astragalin and naringenin. Among the “Talenagao” samples, the three species *R. capitatum*, *R. nivale*, and *R. thymifolium* had average total contents of the 10 compounds of 4.493, 3.665, and 4.372 mg·g^−1^, respectively, which were similar to the 4.705 mg·g^−1^ of the one *R. intricatum* sample. However, the *R. nitidulum* sample had a total content of 2.873 mg·g^−1^, which was much lower than that of the other four species. Among these five species used as “Talenabao,” *R. nivale* had lower quercetin and naringenin contents. *R. thymifolium* had a significantly higher hyperoside content than *R. capitatum* and *R. nivale*, with p-values of 0.000 and 0.004, respectively, and significantly lower contents of guaijaverin and astragalin than *R. capitatum* and *R. nivale*, with p-values of 0.016 and 0.008 for guaijaverin and 0.020 and 0.012 for astragalin, respectively. In addition, *R. intricatum* had high taxifolin and guaijaverin contents and a low avicularin content. However, *R. nitidulum* had a somewhat different compound profile from the other species, with a high astragalin content and low contents of isoquercitrin, avicularin, quercetin, and naringenin.

**Table 5.**
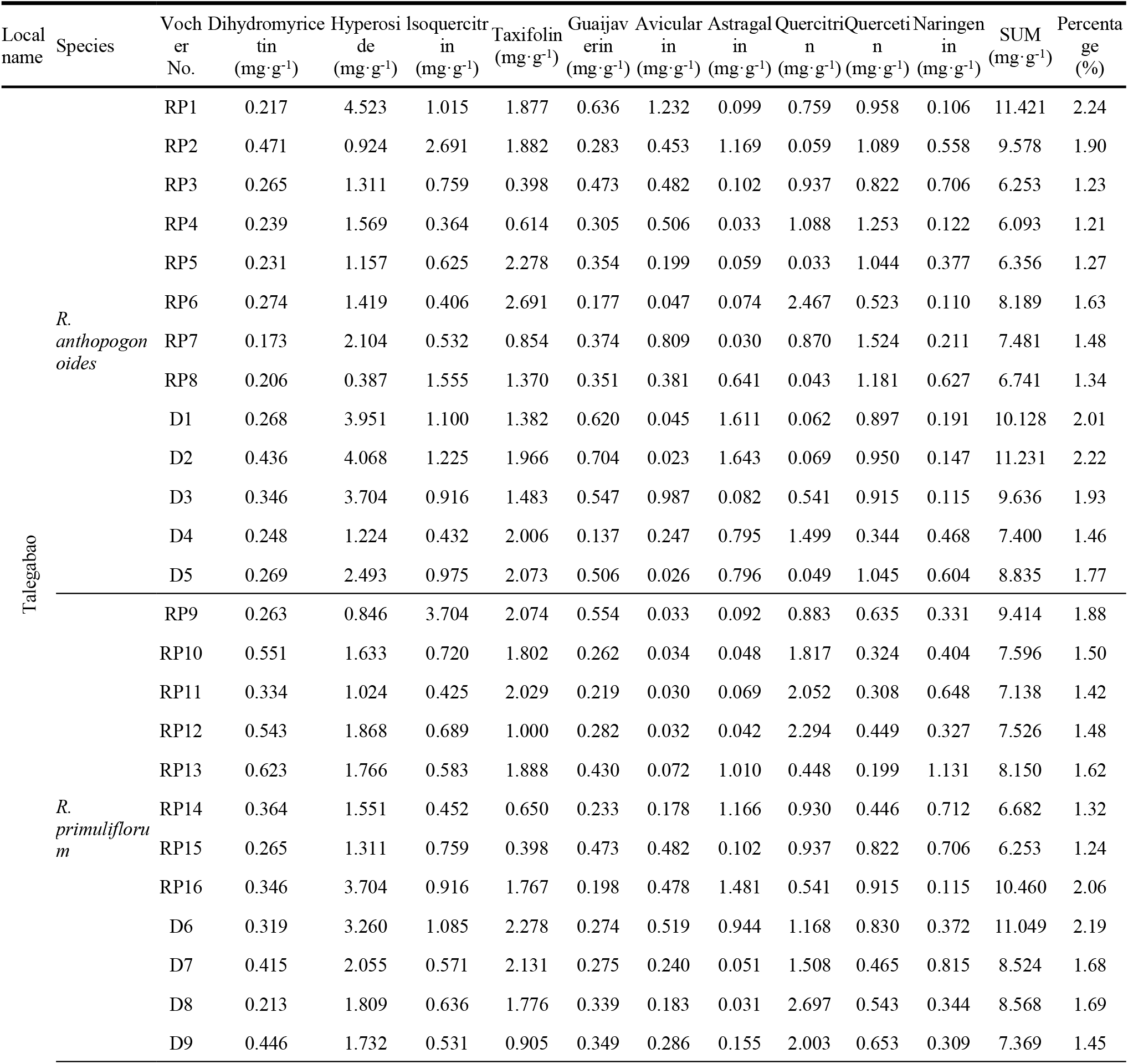
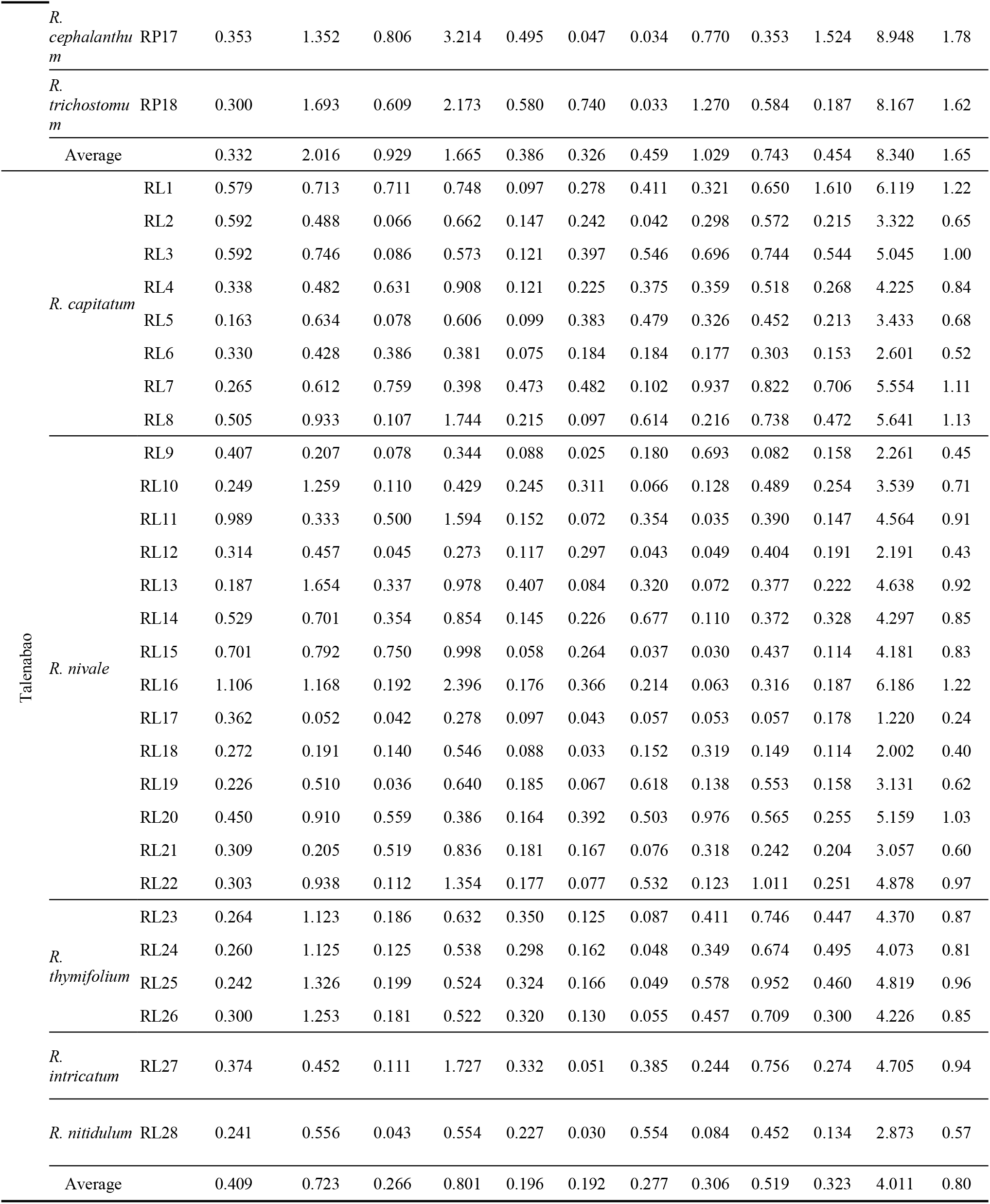
The contents of ten compounds in “Tale” samples.

| Local name | Species | Vocher No. | Dihydromyricetin (mg·g<sup>-1</sup>) | Hyperoside (mg·g<sup>-1</sup>) | Isoquercitrin (mg·g<sup>-1</sup>) | Taxifolin (mg·g<sup>-1</sup>) | Guaijaverin (mg·g<sup>-1</sup>) | Avicularin (mg·g<sup>-1</sup>) | Astragalin (mg·g<sup>-1</sup>) | Quercitrin (mg·g<sup>-1</sup>) | Quercetin (mg·g<sup>-1</sup>) | Naringenin (mg·g<sup>-1</sup>) | SUM (mg·g<sup>-1</sup>) | Percentage (%) |
|---|---|---|---|---|---|---|---|---|---|---|---|---|---|---|
| Talegabao | *R. anthopogonoides* | RP1 | 0.217 | 4.523 | 1.015 | 1.877 | 0.636 | 1.232 | 0.099 | 0.759 | 0.958 | 0.106 | 11.421 | 2.24 |
|  |  | RP2 | 0.471 | 0.924 | 2.691 | 1.882 | 0.283 | 0.453 | 1.169 | 0.059 | 1.089 | 0.558 | 9.578 | 1.90 |
|  |  | RP3 | 0.265 | 1.311 | 0.759 | 0.398 | 0.473 | 0.482 | 0.102 | 0.937 | 0.822 | 0.706 | 6.253 | 1.23 |
|  |  | RP4 | 0.239 | 1.569 | 0.364 | 0.614 | 0.305 | 0.506 | 0.033 | 1.088 | 1.253 | 0.122 | 6.093 | 1.21 |
|  |  | RP5 | 0.231 | 1.157 | 0.625 | 2.278 | 0.354 | 0.199 | 0.059 | 0.033 | 1.044 | 0.377 | 6.356 | 1.27 |
|  |  | RP6 | 0.274 | 1.419 | 0.406 | 2.691 | 0.177 | 0.047 | 0.074 | 2.467 | 0.523 | 0.110 | 8.189 | 1.63 |
|  |  | RP7 | 0.173 | 2.104 | 0.532 | 0.854 | 0.374 | 0.809 | 0.030 | 0.870 | 1.524 | 0.211 | 7.481 | 1.48 |
|  |  | RP8 | 0.206 | 0.387 | 1.555 | 1.370 | 0.351 | 0.381 | 0.641 | 0.043 | 1.181 | 0.627 | 6.741 | 1.34 |
|  |  | D1 | 0.268 | 3.951 | 1.100 | 1.382 | 0.620 | 0.045 | 1.611 | 0.062 | 0.897 | 0.191 | 10.128 | 2.01 |
|  |  | D2 | 0.436 | 4.068 | 1.225 | 1.966 | 0.704 | 0.023 | 1.643 | 0.069 | 0.950 | 0.147 | 11.231 | 2.22 |
|  |  | D3 | 0.346 | 3.704 | 0.916 | 1.483 | 0.547 | 0.987 | 0.082 | 0.541 | 0.915 | 0.115 | 9.636 | 1.93 |
|  |  | D4 | 0.248 | 1.224 | 0.432 | 2.006 | 0.137 | 0.247 | 0.795 | 1.499 | 0.344 | 0.468 | 7.400 | 1.46 |
|  |  | D5 | 0.269 | 2.493 | 0.975 | 2.073 | 0.506 | 0.026 | 0.796 | 0.049 | 1.045 | 0.604 | 8.835 | 1.77 |
|  | *R. primuliflorum* | RP9 | 0.263 | 0.846 | 3.704 | 2.074 | 0.554 | 0.033 | 0.092 | 0.883 | 0.635 | 0.331 | 9.414 | 1.88 |
|  |  | RP10 | 0.551 | 1.633 | 0.720 | 1.802 | 0.262 | 0.034 | 0.048 | 1.817 | 0.324 | 0.404 | 7.596 | 1.50 |
|  |  | RP11 | 0.334 | 1.024 | 0.425 | 2.029 | 0.219 | 0.030 | 0.069 | 2.052 | 0.308 | 0.648 | 7.138 | 1.42 |
|  |  | RP12 | 0.543 | 1.868 | 0.689 | 1.000 | 0.282 | 0.032 | 0.042 | 2.294 | 0.449 | 0.327 | 7.526 | 1.48 |
|  |  | RP13 | 0.623 | 1.766 | 0.583 | 1.888 | 0.430 | 0.072 | 1.010 | 0.448 | 0.199 | 1.131 | 8.150 | 1.62 |
|  |  | RP14 | 0.364 | 1.551 | 0.452 | 0.650 | 0.233 | 0.178 | 1.166 | 0.930 | 0.446 | 0.712 | 6.682 | 1.32 |
|  |  | RP15 | 0.265 | 1.311 | 0.759 | 0.398 | 0.473 | 0.482 | 0.102 | 0.937 | 0.822 | 0.706 | 6.253 | 1.24 |
|  |  | RP16 | 0.346 | 3.704 | 0.916 | 1.767 | 0.198 | 0.478 | 1.481 | 0.541 | 0.915 | 0.115 | 10.460 | 2.06 |
|  |  | D6 | 0.319 | 3.260 | 1.085 | 2.278 | 0.274 | 0.519 | 0.944 | 1.168 | 0.830 | 0.372 | 11.049 | 2.19 |
|  |  | D7 | 0.415 | 2.055 | 0.571 | 2.131 | 0.275 | 0.240 | 0.051 | 1.508 | 0.465 | 0.815 | 8.524 | 1.68 |
|  |  | D8 | 0.213 | 1.809 | 0.636 | 1.776 | 0.339 | 0.183 | 0.031 | 2.697 | 0.543 | 0.344 | 8.568 | 1.69 |
|  |  | D9 | 0.446 | 1.732 | 0.531 | 0.905 | 0.349 | 0.286 | 0.155 | 2.003 | 0.653 | 0.309 | 7.369 | 1.45 |
|  | *R. cephalanthum* | RP17 | 0.353 | 1.352 | 0.806 | 3.214 | 0.495 | 0.047 | 0.034 | 0.770 | 0.353 | 1.524 | 8.948 | 1.78 |
|  | *R. trichostomum* | RP18 | 0.300 | 1.693 | 0.609 | 2.173 | 0.580 | 0.740 | 0.033 | 1.270 | 0.584 | 0.187 | 8.167 | 1.62 |
|  | Average |  | 0.332 | 2.016 | 0.929 | 1.665 | 0.386 | 0.326 | 0.459 | 1.029 | 0.743 | 0.454 | 8.340 | 1.65 |
| Talenabao | *R. capitatum* | RL1 | 0.579 | 0.713 | 0.711 | 0.748 | 0.097 | 0.278 | 0.411 | 0.321 | 0.650 | 1.610 | 6.119 | 1.22 |
|  |  | RL2 | 0.592 | 0.488 | 0.066 | 0.662 | 0.147 | 0.242 | 0.042 | 0.298 | 0.572 | 0.215 | 3.322 | 0.65 |
|  |  | RL3 | 0.592 | 0.746 | 0.086 | 0.573 | 0.121 | 0.397 | 0.546 | 0.696 | 0.744 | 0.544 | 5.045 | 1.00 |
|  |  | RL4 | 0.338 | 0.482 | 0.631 | 0.908 | 0.121 | 0.225 | 0.375 | 0.359 | 0.518 | 0.268 | 4.225 | 0.84 |
|  |  | RL5 | 0.163 | 0.634 | 0.078 | 0.606 | 0.099 | 0.383 | 0.479 | 0.326 | 0.452 | 0.213 | 3.433 | 0.68 |
|  |  | RL6 | 0.330 | 0.428 | 0.386 | 0.381 | 0.075 | 0.184 | 0.184 | 0.177 | 0.303 | 0.153 | 2.601 | 0.52 |
|  |  | RL7 | 0.265 | 0.612 | 0.759 | 0.398 | 0.473 | 0.482 | 0.102 | 0.937 | 0.822 | 0.706 | 5.554 | 1.11 |
|  |  | RL8 | 0.505 | 0.933 | 0.107 | 1.744 | 0.215 | 0.097 | 0.614 | 0.216 | 0.738 | 0.472 | 5.641 | 1.13 |
|  | *R. nivale* | RL9 | 0.407 | 0.207 | 0.078 | 0.344 | 0.088 | 0.025 | 0.180 | 0.693 | 0.082 | 0.158 | 2.261 | 0.45 |
|  |  | RL10 | 0.249 | 1.259 | 0.110 | 0.429 | 0.245 | 0.311 | 0.066 | 0.128 | 0.489 | 0.254 | 3.539 | 0.71 |
|  |  | RL11 | 0.989 | 0.333 | 0.500 | 1.594 | 0.152 | 0.072 | 0.354 | 0.035 | 0.390 | 0.147 | 4.564 | 0.91 |
|  |  | RL12 | 0.314 | 0.457 | 0.045 | 0.273 | 0.117 | 0.297 | 0.043 | 0.049 | 0.404 | 0.191 | 2.191 | 0.43 |
|  |  | RL13 | 0.187 | 1.654 | 0.337 | 0.978 | 0.407 | 0.084 | 0.320 | 0.072 | 0.377 | 0.222 | 4.638 | 0.92 |
|  |  | RL14 | 0.529 | 0.701 | 0.354 | 0.854 | 0.145 | 0.226 | 0.677 | 0.110 | 0.372 | 0.328 | 4.297 | 0.85 |
|  |  | RL15 | 0.701 | 0.792 | 0.750 | 0.998 | 0.058 | 0.264 | 0.037 | 0.030 | 0.437 | 0.114 | 4.181 | 0.83 |
|  |  | RL16 | 1.106 | 1.168 | 0.192 | 2.396 | 0.176 | 0.366 | 0.214 | 0.063 | 0.316 | 0.187 | 6.186 | 1.22 |
|  |  | RL17 | 0.362 | 0.052 | 0.042 | 0.278 | 0.097 | 0.043 | 0.057 | 0.053 | 0.057 | 0.178 | 1.220 | 0.24 |
|  |  | RL18 | 0.272 | 0.191 | 0.140 | 0.546 | 0.088 | 0.033 | 0.152 | 0.319 | 0.149 | 0.114 | 2.002 | 0.40 |
|  |  | RL19 | 0.226 | 0.510 | 0.036 | 0.640 | 0.185 | 0.067 | 0.618 | 0.138 | 0.553 | 0.158 | 3.131 | 0.62 |
|  |  | RL20 | 0.450 | 0.910 | 0.559 | 0.386 | 0.164 | 0.392 | 0.503 | 0.976 | 0.565 | 0.255 | 5.159 | 1.03 |
|  |  | RL21 | 0.309 | 0.205 | 0.519 | 0.836 | 0.181 | 0.167 | 0.076 | 0.318 | 0.242 | 0.204 | 3.057 | 0.60 |
|  |  | RL22 | 0.303 | 0.938 | 0.112 | 1.354 | 0.177 | 0.077 | 0.532 | 0.123 | 1.011 | 0.251 | 4.878 | 0.97 |
|  | *R. thymifolium* | RL23 | 0.264 | 1.123 | 0.186 | 0.632 | 0.350 | 0.125 | 0.087 | 0.411 | 0.746 | 0.447 | 4.370 | 0.87 |
|  |  | RL24 | 0.260 | 1.125 | 0.125 | 0.538 | 0.298 | 0.162 | 0.048 | 0.349 | 0.674 | 0.495 | 4.073 | 0.81 |
|  |  | RL25 | 0.242 | 1.326 | 0.199 | 0.524 | 0.324 | 0.166 | 0.049 | 0.578 | 0.952 | 0.460 | 4.819 | 0.96 |
|  |  | RL26 | 0.300 | 1.253 | 0.181 | 0.522 | 0.320 | 0.130 | 0.055 | 0.457 | 0.709 | 0.300 | 4.226 | 0.85 |
|  | *R. intricatum* | RL27 | 0.374 | 0.452 | 0.111 | 1.727 | 0.332 | 0.051 | 0.385 | 0.244 | 0.756 | 0.274 | 4.705 | 0.94 |
|  | *R. nitidulum* | RL28 | 0.241 | 0.556 | 0.043 | 0.554 | 0.227 | 0.030 | 0.554 | 0.084 | 0.452 | 0.134 | 2.873 | 0.57 |
|  | Average |  | 0.409 | 0.723 | 0.266 | 0.801 | 0.196 | 0.192 | 0.277 | 0.306 | 0.519 | 0.323 | 4.011 | 0.80 |

**Fig 2.**
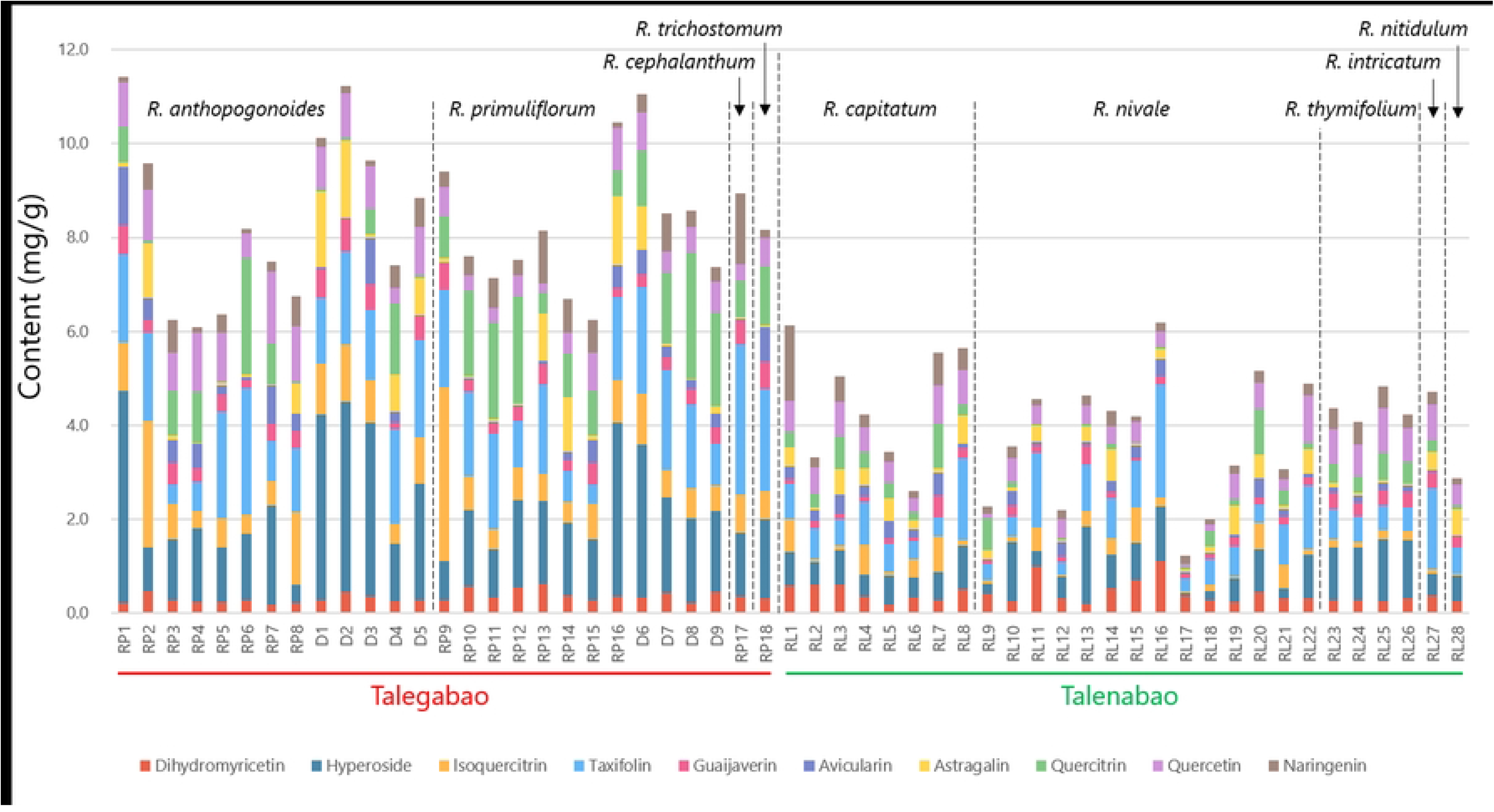
The contents of ten compounds in “Tale” samples.

OPLS-DA was conducted using HPLC data from all 55 *Rhododendron* samples. The OPLS-DA score and loading plots are shown in Fig 3. In the score plot, the “Talegabao” samples were distinguished from the “Talenabao” samples. However, samples within each group were not clearly separated, indicating similar chemical profiles (Fig 3a). In the loading plot, the five species used as “Talenabao” clustered together and were associated with a high dihyromyricetin content. The four species used as “Talegabao” clustered in the right part of the plot but were separated from one another. *R. anthopogonoides* was located in the upper right, distant from the center, and was associated with high quercetin and avicularin content. *R. primuliflorum* and *R. trichostomum* were located in the lower left and were associated with high contents of naringenin, quercitrin, and taxifolin. In addition, *R. cephalanthum* was associated with a high astragalin content (Fig 3b). These OPLS-DA results were consistent with the HPLC analysis.

**Fig 3.**
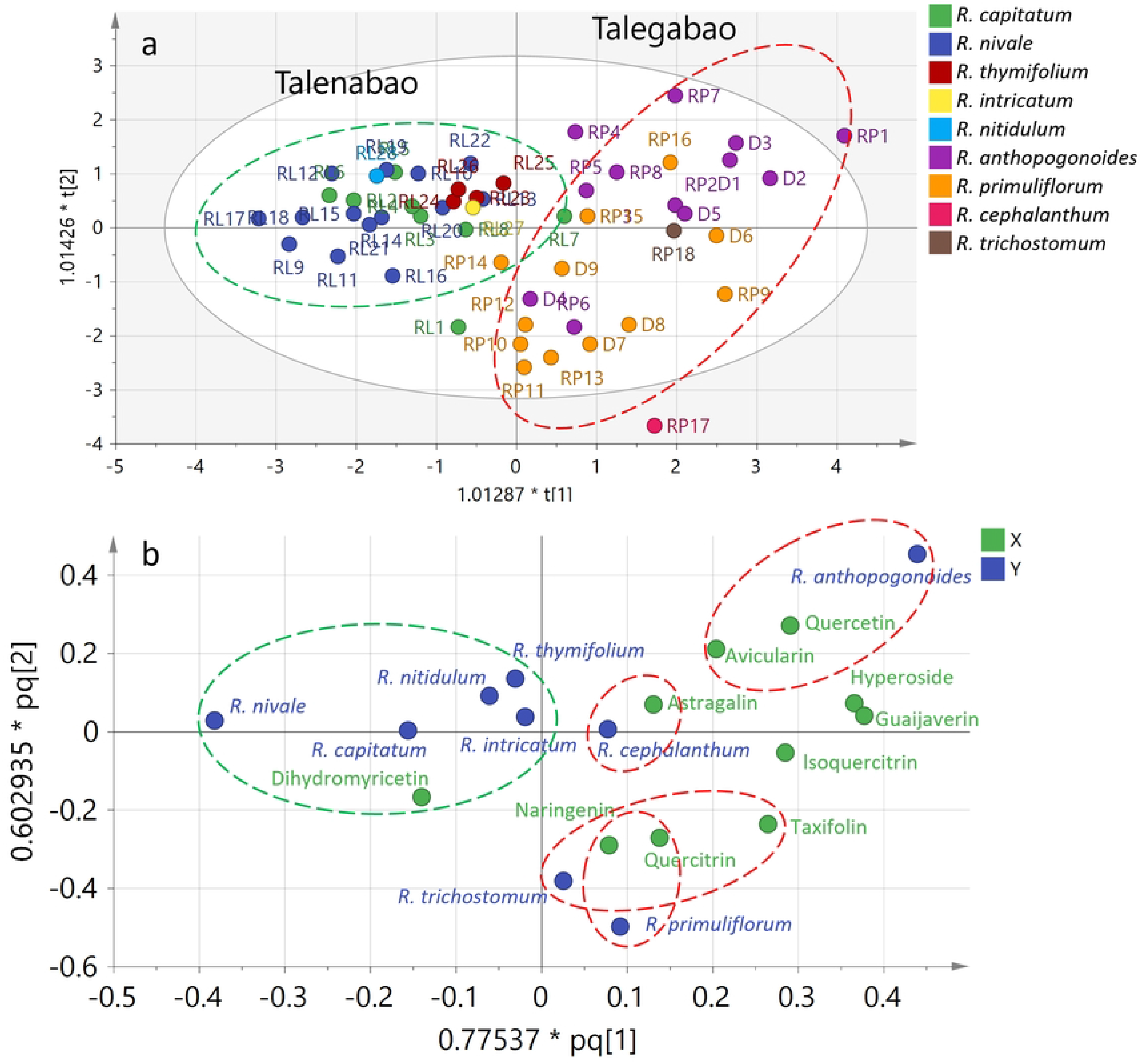
OPLS-DA results of 55 “Tale” samples using the data matrix created by HPLC quantitative data. a, score plot of OPLS-DA results; b, loading plot of OPLS-DA results.

## Discussion

### Molecular identification of *Rhododendron* species

In this study, the ITS2 and *psb*A–*trn*H regions of all nine species used as the traditional Tibetan medicine “Tale” were amplified and sequenced to distinguish among the species. The alignment results indicated that the ITS2 region had limited discriminatory power for the *Rhododendron* species used as “Tale” medicines. In contrast, the *psb*A–*trn*H region successfully distinguished the species used as “Talegabao” and “Talenabao” and showed a high capability for discriminating *R. anthopogonoides* and *R. primuliflorum* from *R. cephalanthum* and *R. trichostomum*. These results are consistent with the findings of Yan et al., who reported that *psb*A–*trn*H had relatively higher species-discrimination power than ITS sequences and recommended it as a single DNA barcode for *Rhododendron* species [14]. However, the species-discrimination power of the DNA barcodes evaluated in the present study was relatively low. Neither ITS2 nor *psb*A–*trn*H could distinguish all nine species from one another.

Interestingly, some studies have reported high discriminatory power of the ITS region for classifying *Rhododendron* species. Gao Lian-Ming et al. reconstructed a phylogenetic tree based on ITS sequences from 12 species representing two sections of Subgenus *Tsutsusi* and four species representing three Sections of Subgenus *Pentanthera* of *Rhododendron*. The strict-consensus tree topology showed that Subgenus *Tsutsusi* was monophyletic, and *R. tashiroi* should be a member of Section *Brachycalyx*, whereas *R. tsusiophyllum* should be a member of Section *Tsutsusi*. ITS sequences successfully resolved the taxonomic relationships of species in Subgenera *Tsutsusi* and *Pentanthera* [16]. Cai Hong-Yue et al. evaluated the relationships among species in the subgenera of *Rhododendron* and clarified the classification of five native *Rhododendron* species in Mount Tai based on ITS regions. The results showed that, among the five species from Mount Tai, *R. mariesii* did not cluster within the branch of Subgenus *Tsutsusi*, whereas the other four species were located in the major branches of their subgenera [17].

However, the species examined in these studies were mainly distributed at lower elevations. The species in the present study were collected from the Qinghai–Xizang Plateau at an average altitude of more than 4,000 m, similar to the species examined by Yan et al., which were collected from the Himalaya–Hengduan Mountains. Highland species appear to show lower levels of species discrimination, with shared sequence identities across species and less distinct species boundaries. A similar phenomenon has also been reported in highland species of subsection Ledum in North America [18]. Yan et al. proposed that these closely related highland species may have diverged morphologically in response to strong selection pressures associated with different habitats while undergoing relatively little genetic divergence, possibly because of long generation times and relatively recent speciation and diversification. In addition, incongruent signals between nuclear and plastid gene regions are common in these species and may result from natural hybridization and gene introgression. These factors may collectively contribute to the limited effectiveness of DNA barcoding in distinguishing these species [14].

To address this issue in highland *Rhododendron* species, Fu et al. introduced a genome-skimming method based on the chloroplast genome to distinguish *Rhododendron* species from the Himalayan–Hengduan Mountains. Compared with the 33% of species distinguished using ITS + matK + rbcL + *trn*H–*psb*A, 55% of species were distinguished using plastid genome and nrDNA sequences [19]. Given the limited taxonomic information available for these highland *Rhododendron* species, the *psb*A–*trn*H region currently showed the highest species-discrimination power among the regions evaluated in the present study and has potential for use in distinguishing the traditional Tibetan medicine “Tale.” With increasing attention to the medicinal, environmental, and horticultural value of these *Rhododendron* species, further exploration of new DNA barcode regions and other methods for species discrimination is warranted.

### Quantitative analysis of the 10 compounds in “Tale”

In this study, we conducted a phytochemical investigation of nine *Rhododendron* species used as traditional Tibetan medicine “Tale.” An HPLC method was used to simultaneously detect 10 flavonoid compounds, and the contents of these compounds were determined in the nine *Rhododendron* species. All 10 compounds were detected in the plant specimens and crude drug samples, indicating a high degree of phytochemical similarity among the samples. The nine species used as “Tale” medicine are reasonable. The species used as “Tale” medicine differed primarily in the contents of these compounds. The total content of the 10 flavonoid compounds in “Talegabao” was more than twice that in “Talenabao,” indicating a substantial difference between the two types of medicines. Flavonoid compounds have been associated with various pharmacological activities, including antioxidant, anti-inflammatory, antimicrobial, antitumor, hypoglycemic, hepatoprotective, neuroprotective, and cardioprotective effects [20]. As described in the Introduction, “Talegabao” was described as the first choice among “Tale” medicines, whereas “Talenabao” is usually used as an alternative. Our results provide phytochemical evidence relevant to the use of these medicines.

Among the 10 compounds, nine had higher contents in the species used as “Talegabao,” and only dihydromyricetin had a higher content in the species used as “Talenabao.” Furthermore, the contents of individual compounds varied among species. Different flavonoid compounds have different properties [21]. These differences may contribute to differences in the effects of the two types of medicine. Although the content of certain compounds differed among species, samples within each medicine type had similar total contents and phytochemical profiles, and the OPLS-DA results indicated that species within each type were not clearly distinguishable. These findings support the use of multiple species as sources of “Tale” medicines. “Talenabao” can be used as a substitute for “Talegabao.” Among the species analyzed as “Talegabao” or “Talenabao,” certain compounds showed distinct contents among species, providing a basis for further standardization of “Tale” medicines.

Few studies on the chemical composition of *Rhododendron* species used as “Tale” have been reported. To our knowledge, this study is the first conducted to compare *Rhododendron* species used as “Tale” based on molecular and phytochemical analyses. However, several issues related to the standardization of “Tale” medicines during their production and use remain to be addressed. Multiple species are used as “Tale” medicines, and the contents of individual compounds vary among these species. Differences were observed not only among species but also within species, with some individuals showing distinct compound contents. However, the factors influencing the accumulation of these compounds remain unknown. Although several factors influencing the accumulation of flavonoid compounds have been examined, Maxime assessed the effect of anthropogenic disturbances on the chemical properties of *R. groenlandicum* in the territories of three Indigenous communities. The results showed that flavonoid concentrations varied mainly by territory, whereas disturbance type had a smaller effect [22]. The mechanisms underlying flavonoid accumulation remain unclear.

Multiple monoterpenoid and sesquiterpenoid compounds have been isolated from the essential oils of *Rhododendron* species, and these essential oils have shown multiple properties [23]. In addition, the toxic constituents of *Rhododendron* species warrant attention [2,24]. Future standardization studies should consider additional species and evaluation indicators, including pharmacological studies, essential oils, and toxic compounds.

## Conclusions

In this study, 46 plant specimens and nine crude drug samples belonging to nine *Rhododendron* species were collected from the Qinghai–Xizang Plateau. Molecular methods based on the ITS2 region of nuclear DNA and the *psb*A–*trn*H region of chloroplast DNA were applied to these plant specimens. In addition, 10 flavonoid compounds were quantitatively determined by HPLC. The molecular analysis revealed that the *psb*A–*trn*H region successfully distinguished the species used as “Talegabao” or “Talenabao”; however, ITS2 had limited discriminatory power because of its high degree of conservation. The HPLC analysis showed that all 10 compounds were detected in all samples. The species used as “Talegabao” had more than twice the total content of the 10 flavonoid compounds observed in the species used as “Talenabao.” In addition, differences in the contents of certain compounds were detected among species, which may provide potential chemical markers for the quality evaluation of “Tale” medicines. According to the results, “Talegabao” is the first choice among “Tale” medicines, whereas “Talenabao” can be used as an alternative to “Talegabao”; however, the appropriate dose requires further discussion in clinical practice. The *psb*A–*trn*H region may serve as a DNA barcode for distinguishing the species used as “Talegabao” and “Talenabao.”

## References

1. Hussain A, Azam S, Maqsood R, Anwar R, Akash MSH, Hussain H, et al. Chemistry, biosynthesis, and theranostics of antioxidant flavonoids and polyphenolics of genus *Rhododendron*: an overview. Naunyn-Schmiedeberg’s Arch Pharmacol. 2025;398: 1171–1214. doi:10.1007/s00210-024-03428-6

2. Popescu R, Kopp B. The genus *Rhododendron*: An ethnopharmacological and toxicological review. Journal of Ethnopharmacology. 2013;147: 42–62. doi:10.1016/j.jep.2013.02.022

3. Wu Z, Raven PH, Hong D. Flora of China. Science Press & Missouri Botanical Garden Press; 2005.

4. Yang Y. Zang yao zhi [= Dictionary of traditional Tibetan medicine]. Qinghai People’s Publishing House; 1991.

5. Jiang Q, Bai Y. Research progress of Mongolian drug Rhododendron and plants of the genus Rhododendron. 2014;29: 68–70. doi:10.14045/j.cnki.15-1220.2014.01.015

6. Wang S, Lu Y, Zhang K, Wang S, Zhang S, Su H, et al. The chemical constituents and anti-complement activity of seven *Rhododendron* species in Tibetan medicine. Molecules. 2026;31: 2257. doi:10.3390/molecules31132257

7. Zhuoji Z, Li L, Wang X, Jixian C, Gazangcuo, Gengzangjia. Research advances on chemical constituents, pharmacological effects, and clinical applications of Liexiang Dujuan (*Rhododendron anthopogonoides* Maxim.) in Tibetan medicine. 2005;31: 100–107. doi:10.13862/j.cn43-1446/r.2025.08.016

8. Zhang J, Ma J, Ynag Y, Yao J, Huang L, Gao L, et al. Analysis of essential oil in *Rhododendron anthopogoncides*. 2003;34: 304–305.

9. J H, X S, L D, X Y, B L, Y W, et al. Chemical constituents, antibacterial, acaricidal and anti-inflammatory activities of the essential oils from four *Rhododendron* species. Frontiers in veterinary science. 2022;9. doi:10.3389/fvets.2022.882060

10. Q S, Tt L, Ym W, Xy S, C L, Jy L, et al. Meroterpenoids with diverse structures and anti-inflammatory activities from *Rhododendron anthopogonoides*. Phytochemistry. 2020;180. doi:10.1016/j.phytochem.2020.112524

11. Yan L, Hu F, Wu J, Han F. The biological activity of essential oil in *Rhododendron anthopogonoides* Maxim and *Artemisia dubia* Wall. ex Bess .to grassland caterpillar, Gynaephora menyuanenis ( Lepidoptera :Lymantriidae). 2009;18: 58–63. doi:10.3969/j.issn.1004-1389.2009.05.013

12. K Y, Yx Z, Cf W, Ss D, Zw D, Qz L, et al. Toxicity of *Rhododendron anthopogonoides* essential oil and its constituent compounds towards *Sitophilus zeamais*. Molecules (Basel, Switzerland). 2011;16. doi:10.3390/molecules16097320

13. Liu Y, Zhang L, Chen S, Chen K. Identification of 10 medicinally used *Rhododendron* species based on ITS2 sequences. 2011;34: 1342–1345. doi:10.13863/j.issn1001-4454.2011.09.010

14. Yan L, Liu J, Möller M, Zhang L, Zhang X, Li D, et al. DNA barcoding of *Rhododendron* (Ericaceae), the largest Chinese plant genus in biodiversity hotspots of the Himalaya–Hengduan Mountains. Molecular Ecology Resources. 2015;15: 932–944. doi:10.1111/1755-0998.12353

15. Liu Q, Yu H, Dong Y, Quan W, Su Z, Li L. Quality evaluation of Lonicerae Flos produced in Southwest China based on HPLC analysis and antioxidant activity. Molecules. 2024;29: 2560. doi:10.3390/molecules29112560

16. Gao L, Yang J, Zhang C, Li D. Phylogenetic relationship of subgenus *Tsutsusi* ( *Rhododendron*) based on ITS sequences. 2002;24: 313–320.

17. Cai H, Xing S, He R, Zhou B, Zhao F. Phylogenetic relationship analysis of genus *Rhododendron* (Ericaceae) from Mountain Tai based on ITS2 sequences and morphological traits. 2017;53: 1489–1498. doi:10.13592/j.cnki.ppj.2017.0002

18. Iunusova DR, Polezhaeva MA, Marchuk EA. Confusing boundaries of the labrador tea species: dispersal history explains the lack of clear species structure. Annals of Botany. 2024;134: 959–974. doi:10.1093/aob/mcae130

19. Fu C, Mo Z, Yang J, Cai J, Ye L, Zou J, et al. Testing genome skimming for species discrimination in the large and taxonomically difficult genus *Rhododendron*. Molecular Ecology Resources. 2022;22: 404–414. doi:10.1111/1755-0998.13479

20. M V, L R. Phytochemical profiling and biological activities of *Rhododendron* subsect. *Ledum*: discovering the medicinal potential of labrador tea species in the northern hemisphere. Plants (Basel, Switzerland). 2024;13. doi:10.3390/plants13060901

21. Wang H, Li Y, Shen W, Guo G, Jiang Z, Cen Y, et al. Studies on antiproliferative effect of flavones compounds Isolated from Yao herb medicines. 2007;30: 980–984. doi:10.13863/j.issn1001-4454.2007.08.005

22. Thomas M, Lamara M, Asselin H, Fenton NJ. Effects of industrial disturbances on the flavonoid concentration of *Rhododendron groenlandicum*. Botany. 2023;101: 343–356. doi:10.1139/cjb-2022-0136

23. Y S, P W, J L, Y H, X W. Chemical composition of essential oils from three *Rhododendron* species and their repellent, insecticidal and fumigant activities. Chemistry & biodiversity. 2022;19. doi:10.1002/cbdv.202200740

24. Q L, Jx H, Xm Z, Wh X. Traditional uses, phytochemistry, pharmacology, toxicology, and quality control of *Rhododendron dauricum* L. leaves: A comprehensive review. Journal of ethnopharmacology. 2023;305. doi:10.1016/j.jep.2022.116085

